# Comprehensive and accurate genetic variant identification from contaminated and low coverage *Mycobacterium tuberculosis* whole genome sequencing data

**DOI:** 10.1101/2021.09.16.460612

**Authors:** Tim H. Heupink, Lennert Verboven, Robin M. Warren, Annelies Van Rie

## Abstract

Improved understanding of the genomic variants that allow *Mycobacterium tuberculosis* (*Mtb*) to acquire drug resistance, or tolerance, and increase its virulence are important factors in controlling the current tuberculosis epidemic. Current approaches to *Mtb* sequencing however cannot reveal *Mtb*’s full genomic diversity due to the strict requirements of low contamination levels, high *Mtb* sequence coverage, and elimination of complex regions.

We developed the XBS (compleX Bacterial Samples) bioinformatics pipeline which implements joint calling and machine-learning-based variant filtering tools to specifically improve variant detection in the important *Mtb* samples that do not meet these criteria, such as those from unbiased sputum samples. Using novel simulated datasets, that permit exact accuracy verification, XBS was compared to the UVP and MTBseq pipelines. Accuracy statistics showed that all three pipelines performed equally well for sequence data that resemble those obtained from high depth coverage and low-level contamination culture isolates. In the complex genomic regions however, XBS accurately identified 9.0% more single nucleotide polymorphisms and 8.1% more single nucleotide insertions and deletions than the WHO-endorsed unified analysis variant pipeline. XBS also had superior accuracy for sequence data that resemble those obtained directly from sputum samples, where depth of coverage is typically very low and contamination levels are high. XBS was the only pipeline not affected by low depth of coverage (5-10×), type of contamination and excessive contamination levels (>50%). Simulation results were confirmed using WGS data from clinical samples, confirming the superior performance of XBS with a higher sensitivity (98.8%) when analysing culture isolates and identification of 13.9% more variable sites in WGS data from sputum samples as compared to MTBseq, without evidence for false positive variants when ribosomal RNA regions were excluded.

The XBS pipeline facilitates sequencing of less-than-perfect *Mtb* samples. These advances will benefit future clinical applications of *Mtb* sequencing, especially whole genome sequencing directly from clinical specimens, thereby avoiding *in vitro* biases and making many more samples available for drug resistance and other genomic analyses. The additional genetic resolution and increased sample success rate will improve genome-wide association studies and sequence-based transmission studies.

**Impact statement:** *Mycobacterium tuberculosis* (Mtb) DNA is usually extracted from culture isolates to obtain high quantities of non-contaminated DNA but this process can change the make-up of the bacterial population and is time-consuming. Furthermore, current analytic approaches exclude complex genomic regions where DNA sequences are repeated to avoid inference of false positive genetic variants, which may result in the loss of important genetic information.

We designed the compleX Bacterial Sample (XBS) variant caller to overcome these limitations. XBS employs joint variant calling and machine-learning-based variant filtering to ensure that high quality variants can be inferred from low coverage and highly contaminated genomic sequence data obtained directly from sputum samples. Simulation and clinical data analyses showed that XBS performs better than other pipelines as it can identify more genetic variants and can handle complex (low depth, highly contaminated) Mtb samples. The XBS pipeline was designed to analyse Mtb samples but can easily be adapted to analyse other complex bacterial samples.

**Data summary:** Simulated sequencing data have been deposited in SRA BioProject PRJNA706121. All detailed findings are available in the Supplementary Material. Scripts for running the XBS variant calling core are available on https://github.com/TimHHH/XBS The authors confirm all supporting data, code and protocols have been provided within the article or through supplementary data files.

## INTRODUCTION

Genetic approaches are increasingly used in tuberculosis research and for the diagnosis of drug resistant tuberculosis. Whole genome sequencing (WGS) of *Mycobacterium tuberculosis* (*Mtb*) aims to investigate the entire genome of the *Mtb* strain to comprehensively assess all known drug resistance conferring regions, provide maximum resolution for genetic transmission studies, and investigate the role of genomic variants using genome wide association studies [1]. The three key problems facing the current *Mtb* WGS approaches are the need for high quantities of *Mtb* DNA, presence of contaminant bacterial and human DNA, and the presence of complex regions in the *Mtb* genome.

*Mtb* is notoriously difficult to sequence directly from clinical samples because the DNA from human cells, bacteria and viruses outnumbers that from *Mtb* bacilli. This results in insufficient template *Mtb* DNA and low genomic depth of coverage when sequenced [2]. *Mtb* WGS therefore primarily uses cultured isolates, which requires a harsh decontamination step followed by a lengthy (2 to 4 weeks) incubation to generate high quantities of *Mtb*. The decontamination step not only reduces the presence of bacteria other than *Mtb*, but may also reduce the *Mtb* load [3]. The culture step can increase the presence of certain strains over others due to stochastic processes or when certain strains are better suited at growing in culture media [4,5]. These processing steps thus result in a population bias, where the inferred *in vitro Mtb* population may not truly represent the *in vivo* population. To generate a rapid and unbiased result, *Mtb* would thus ideally be sequenced directly from the clinical sample.

Despite decontamination, a small proportion of contaminants may persist in the DNA extracted from the culture isolate. Current *Mtb* bioinformatic pipelines use *in silico* meta-genomic classification software to identify these contaminants and exclude samples with a high proportion of contaminant DNA. For example the unified analysis variant pipeline (UVP) uses a cut-off of maximum 10% contamination [6], which may exclude valuable samples from analysis. In addition to the low contamination threshold, the *Mtb* community has adopted relatively high standards for genome coverage, with 30× to 50× and up to 100× depth being the most commonly used. A Poisson distribution however reveals that a mean depth of coverage of 15.8× results in 95% of the genome being covered by 10× or more reads (Lander and Waterman 1988), which should be sufficient for accurately calling majority variants in a haploid genome.

A third problem that complicates *Mtb* WGS is the abundance of complex regions including repeats, transposons, duplicates and phage genes, and the numerous PE/PPE genes. These complex regions are generally excluded from analyses by *Mtb* pipelines. The core genome multi-locus sequence typing (cgMLST) method goes even further as only the most trustable regions are analysed by cgMLST [7]. While these strategies ensure the accuracy of the genome assembly and variant calling, they can result in the loss of a significant proportion of genomic information (∼9% when using the UVP pipeline [6]) that may be important for defining transmission events and identification of variants that affect pathogenicity.

There is thus a need for novel bioinformatics tools that overcome the current requirements of low contamination, high *Mtb* DNA sequence coverage and exclusion of complex genomic regions [8]. The Genome Analysis ToolKit (GATK), originally designed for human genome studies (i.e. diploid), now allow for processing of haploid genomes such that of *Mtb*. The Base Quality Score Recalibration and Indel Realigner tools and single sample variant calling using the now superseded Unified Genotyper tool have already been applied in *Mtb* genome studies [6,9]. The GATK’s ‘Germline short variant discovery Best Practices workflow’ however includes joint genotyping and machine-learning-based variant filtering and has seen little to no implementation in bacterial and *Mtb* genome assembly pipelines. The major advantage of joint variant calling, as opposed to single sample variant calling, is a greater sensitivity for variants at low frequency in the population and detection of variants in low coverage samples for which there would be insufficient confidence if the sample had been analysed on its own [10]. Joint variant calling also enables the calculation of various statistical annotations (including depth of coverage, strand bias and read mapping quality) for alleles in the population rather than for those in a single strain. These population variants are more numerous and their annotations suffer from fewer stochastic deviations, thereby improving subsequent variant filtering. The machine-learning-based variant filtering (VQSR) in the GATK [11] eliminates the need for hard-filtering of variants, which is commonly applied through the use of rather arbitrary cut-offs for strand bias and coverage depth.

We hypothesize that the GATK’s tools are suitable for distinguishing contaminant variants from *Mtb* and to score and identify variants in complex regions of the *Mtb* genome. We developed a novel *Mtb* pipeline integrating the GATK’s tools to improve the identification of genetic variants in less-than-perfect *Mtb* samples and thereby greatly increase our power to capture the diversity of within-patient and within-bacterial genetic information. We also tested the variant calling core of this pipeline for its accuracy to identify genetic variants in comparison to existing pipelines.

## METHODS

### Development of the XBS pipeline

The compleX Bacterial Sample (XBS) pipeline was designed to perform analyses of Illumina FASTQ sequence data. The pipeline was primarily designed to analyse Mtb samples but can easily be adapted to analyse other complex bacterial samples. XBS was realised through coupling published software packages with custom Bash and Python scripts.

Pipelines typically start with identifying the level of contaminants and/or removing contaminants before mapping the sequence reads. In XBS, all FASTQ sequence data, whether single read or paired-end, are directly mapped to the reference genome (H37Rv: NC_000962.3) however using BWA mem [12] (Figure 1). XBS does not employ an adapter trimming step because BWA mem locally aligns sequence reads, which masks the portions of the read that do not align well with the reference genome. Skipping the step of removing contaminants saves considerable computing time but does require sophisticated downstream variant filtering to distinguish genuine *Mtb* variants from those introduced by contaminants.

**Figure 1:**
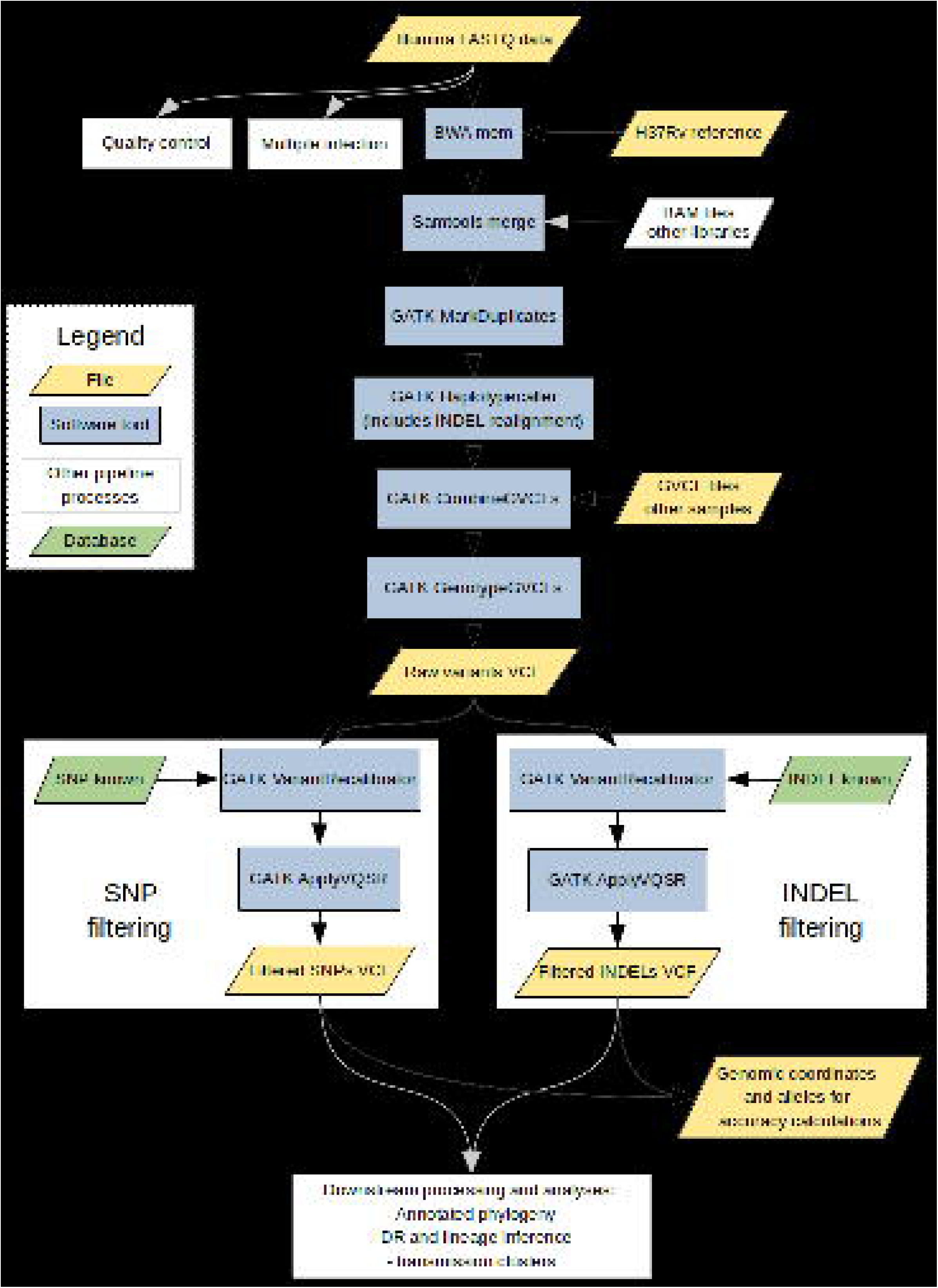
Flow chart for XBS’s variant calling core.

Next, the mapped sequence library is merged with other independently mapped sequence libraries from the same sample using Samtools (https://www.htslib.org/). The GATK MarkDuplicates (Picard) is then used to mark duplicated reads in the merged bam file. Unlike other *Mtb* pipelines XBS does not employ Base Quality Score Recalibration (BQSR) to avoid that variants in contaminant DNA are interpreted as systematic error by BQSR which would result in reduced base qualities, including for genuine *Mtb* variants.

The mapped sequences are then locally reassembled to correctly identify possible haplotypes and their variants using the GATK HaplotypeCaller. At this point, the statistics of depth of coverage, breadth of coverage, multiple infection and level of nontuberculous Mycobacteria (NTM) contamination are assessed to judge if a sample is suitable for subsequent joint variant calling. The coverage statistics are inferred using the GATK CollectWgsMetrics (Picard). Quality approved samples’ Genomic Variant Call Format (GVCF) files are then merged with the GATK CombineGVCFs and the genotypes are joint called using the GATK GenotypeGVCFs. This results in a VCF file with the unfiltered variants for all quality approved samples. GATK is run with a ploidy of 1 for the variant calling processes so that the allele with the highest confidence is identified as the allele representing the haploid genotype for each variant site.

Next, the machine-learning-based variant filtering (VQSR) in the GATK is employed to identify the likely true variants [11]. This step requires a truth set of variants known to occur in *Mtb*, which can for example consist of DR conferring mutations. Single nucleotide polymorphisms (SNPs) and insertions and deletions (INDELs) are processed separately for variant filtering. The annotated statistics calculated during the genotyping are used to build a positive statistical model for the variants in the dataset that also occur in the truth set. Similarly, a negative variant model is built for the variants with the most inferior annotated statistics. The remaining variants not consulted for either the positive or the negative model construction are then confidence scored according to the placement of their annotated statistics in relation to these models. To identify as many variants as possible, variants are then filtered by applying a target sensitivity of 99.9%, calculated as the percentage of identified variants from the present truth-set variants. The filtered SNP and INDEL VCFs are then further processed as appropriate for constructing annotated phylogenies and inferences of multiple infection, drug (hetero-)resistance, lineage and transmission clusters.

### In silico development of a simulated dataset

A dataset of 1,200 simulated samples representing 50,000 SNPs, 2,500 insertions and 2,500 deletions was developed (Figure 2). Of these, 600 were designed to resemble WGS data from mono-culture isolates and 600 to resemble WGS data obtained directly from sputum samples, the latter including high levels of various contaminants.

**Figure 2:**
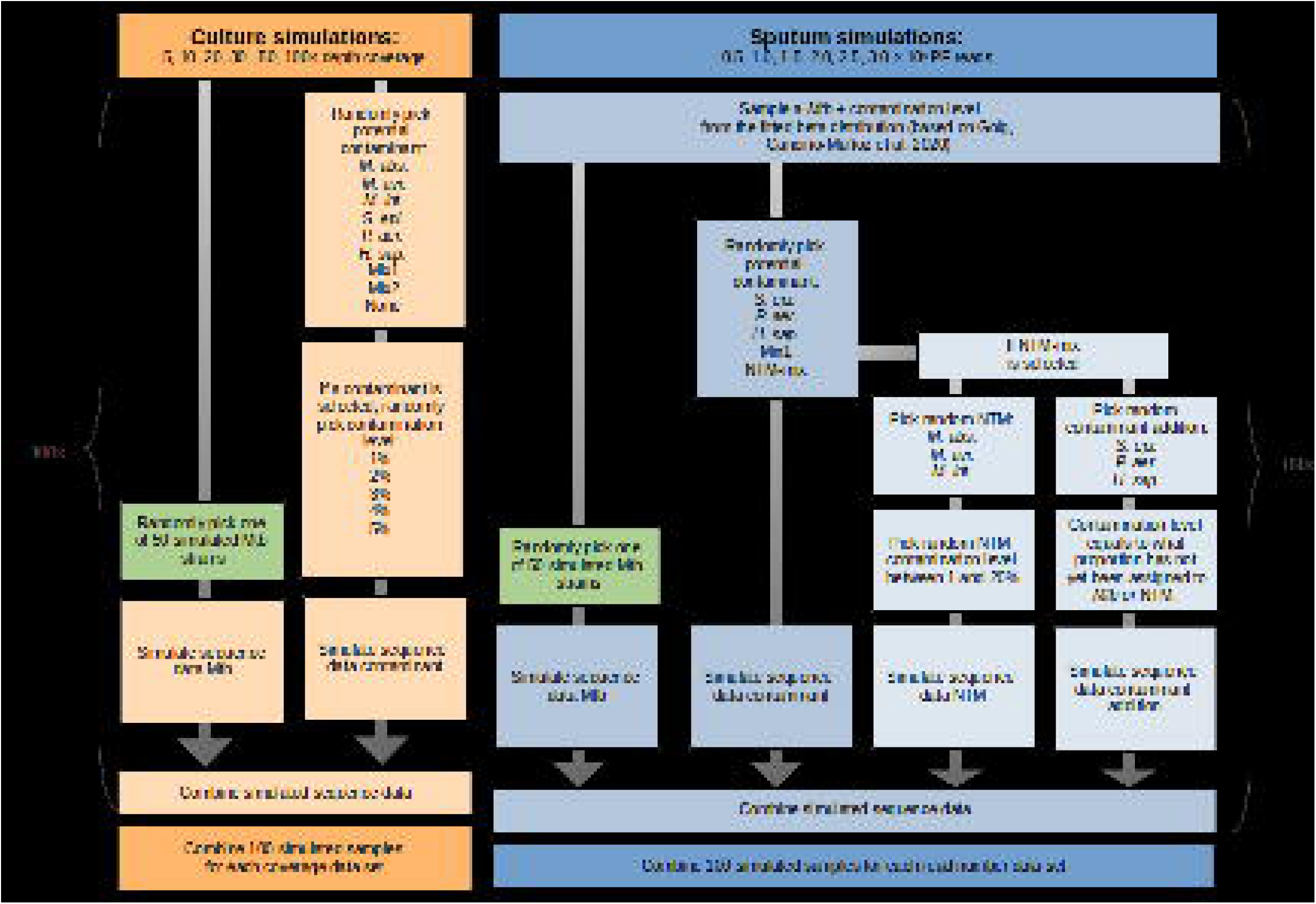
Flow chart for dataset construction.

First, a set of 50 simulated strains with the exact mutations known in respect to the reference genome was created *in silico* using SNP Mutator v.1.2.0 [13]. Each simulated genome was created by randomly introducing 1000 SNPs, 50 single nucleotide insertions and 50 single nucleotide deletions in H37Rv (NC_000962.3). Multi-nucleotide INDELs could not be introduced using the SNP Mutator software. SNPs and INDELs were introduced randomly throughout each simulated strain genome to ensure that random bases were affected and/or introduced and to create genetically varying strains. The GATK LeftAlignAndTrimVariants was used so that the truth set its INDELs were in the standard notation.

For each dataset, 100 strains were randomly drawn from the 50 simulated strains to ensure that some strains and their variants occurred more than once, as would be the case for clinical datasets were specific strains and drug resistance variants often occur more than once. To simulate WGS data obtained from culture isolates, 6 datasets representing 5×, 10×, 20×, 30×, 50× or 100× depth were generated using a 100 randomly drawn simulated strains each (Figure 2). In order to investigate the impact of low-level contamination, no or low-level contamination (0, 1, 2, 3, 4, or 5%) and contamination type was randomly assigned to each of the simulated strains. The eight contamination types used were *Mycobacterium intracellulare* (NC_016946.1), *Mycobacterium abscessus* (NC_010397.1, including plasmid), *Mycobacterium avium* (NC_002944.2), the three most common NTM, *Pseudomonas aeruginosa* (NC_002516.2) and *Staphylococcus epidermidis* (NC_004461.1), *Homo sapiens* (GRCh38), a mixture of the three NTM (*M. intracellulare, M. abcessus* and *M. avium*) and a mixture of all six contaminants. Because it was not possible to represent the full diverse spectrum of contaminating bacteria in the simulations, *Pseudomonas aeruginosa* and *Staphylococcus epidermidis* were selected as these are the most common bacterial contaminants [2], NTMs were included because these pose a serious challenge for Mtb variant calling. The simulated samples included no or low-level contamination in order to resemble WGS data from culture isolates and to be able to investigate the effect of the various low levels of such contamination. Simulated contaminant sequence reads were added to the simulated *Mtb* reads so that the final contamination level matched the assigned contamination percentage.

ART v.2.5.8 software [14] was then used to emulate Illumina 150 bp paired-end sequence reads with an HiSeq error profile and a Poisson distributed 300bp average library insert size for each simulated strain and its contaminant(s). In total, 600 cultured WGS samples were generated in six datasets of 100 simulated samples with each dataset representing a different level of coverage (5×, 10×, 20×, 30×, 50× or 100× depth) to allow assessment of the relation between coverage and accuracy of variant identification.

To simulate *Mtb* WGS data obtained directly from sputum samples, another six datasets were generated, each with a set number of paired-end sequence reads per sample, ranging from 500,000 to 3,000,000 PE reads (Figure 2). *Mtb* and contamination levels were randomly sampled from a beta distribution around 0.01 to 78.63% *Mtb* DNA to match levels observed for direct-from-sputum WGS data [8]. The contamination type was either *P. aeruginosa, S. epidermidis, H. sapiens*, a mixture of *P. aeruginosa, S. epidermidis* and *H. sapiens* or a NTM mixture (*M. intracellulare, M. abcessus* or *M. avium up to 20% of the Mtb fraction with* the remaining contamination consisting of *P. aeruginosa, S. epidermidis* or *H. sapiens*).

ART v.2.5.8 software [14] was used as described previously to emulate sequence reads for each simulated strain and its contaminant(s). In total, 600 simulated WGS data directly from sputum samples were generated in six datasets of 100 simulated samples, each dataset representing a number of paired-end sequence reads and had various levels and types of contamination, allowing the study of the relation between contaminant nature, read number and variant identification.

### Assessment of XBS pipeline performance

The performance of XBS was compared to UVP [6] and MTBseq [15], two well-established and commonly used *Mtb* pipelines. Each pipeline was evaluated using their standard settings. For UVP, variants in the GATK filtered VCF file were used for accuracy calculations. For MTBseq, two approaches were assessed. In ‘MTBseq-basic’, the ‘GATK position variants’ file was used for accuracy calculations. In ‘MTBseq-exrep’, the variants marked ‘repetitive’ in MTBseq’s ‘MTB_Gene_Categories.txt’ were excluded to assess the effect of this commonly applied filtering step. From here on XBS, UVP and the two MTBseq approaches will be referred to as pipelines.

For XBS, VQSR was run with a truth set consisting of 5,000 SNPs, 250 insertions and 250 deletions randomly selected from the mutations known to occur in the 50 simulated strains. The truth set therefore represented 10% (5,500/ 55,000) of the total variants introduced *in silico*. The GATK VariantRecalibrator was run to score each SNP according to the inferred positive and negative models, built on the depth, mapping quality, mapping quality rank-sum and quality by depth statistical annotations. These annotations were processed in an allele specific fashion to distinguish between genuine and contaminant variants occurring on the same genomic location. Annotations that showed insufficient variance, as determined by VQSR, were excluded. A logit transform and jitter were applied to improve mapping quality-based filtering. The FS, ReadPosRanksum and SOR annotations were excluded because they are more applicable for real sequence than for simulated data. Where possible, four Gaussians were used for the positive model. The GATK ApplyVQSR was applied with a truth sensitivity level of 99.9%. The same process was followed for INDELs except that allele specificity was disabled and, where possible, two Gaussians were used for the positive model. The MQ annotation was not taken into consideration following the GATK Best Practices Workflow for Germline short variant discovery. To avoid over-representation of contaminant alleles in the filtered dataset, INDELs in the sputum simulations were filtered using a VQSLOD score of 0 rather than a truth sensitivity level. This ensures that only those INDELs that are most likely to fall in the positive and not the negative model are kept. The resulting SNP and INDEL variants were used for the accuracy calculations. Version 4.1.9.0 of the GATK was used.

The six datasets simulating WGS from culture were analysed using all four pipelines (XBS, MTBseq-basic, MTBseq-exrep, UVP). The six datasets simulating WGS from sputum samples only by three pipelines as UVP can not analyse samples with contamination exceeding 10%. The inferred variants identified by each pipeline were compared with the truth in terms of genome position and allelic nature (bases involved and length). The pipeline’s accuracy was calculated in terms of precision, recall and their harmonic mean (F_1_ score).

For simulation of WGS from culture isolates, accuracy scores were averaged over the 100 samples in each dataset and calculated for each combination of variant type (SNP or INDEL) and genomic region (complete, complex or non-complex). F_1_ scores were plotted for the four pipelines at six levels of depth for the complete genome, and for complex and non-complex regions separately. UVP’s list of excluded loci was used to define regions as complex, these were filtered using the GATK SelectVariants.

For simulation of WGS from sputum, the range of F_1_ scores was calculated, separately for SNPs and INDELs, and the proportion of samples with an F_1_ score >0.9 was estimated. The number of *Mtb* reads was converted to the theoretical depth of coverage (number of simulated *Mtb* reads multiplied by average read length) and plotted against F_1_ scores for each pipeline, contamination level and type after excluding samples with a theoretical coverage of <20× so that the lesser performance of such samples did not distort these figures.

Plots were created in R using the ggplot2 and gridExtra packages.

### Analysis of WGS from clinical samples

The performance of the XBS variant caller was examined using two published WGS datasets obtained from clinical samples and compared to UVP and MTBseq. Data published by Roetzer et al. [16] was used to test the pipelines’ sensitivity by identifying a set of known variants (Sanger confirmed) in DNA extracted from cultured Mtb samples. Data from Goig et al. [8] was used to evaluate the ability to call variants in WGS data from DNA extracted directly from sputum. Only samples with ≥5× genomic coverage depth (S02, S26, S17, S21, S31, S20, S27, S67, S09 and S69) were analysed to ensure sufficient width of coverage and prevent problems with phylogenetic inference. A reference set of 125 diverse cultured Mtb strains with high coverage WGS data was included in the analyses to provide a reference in the phylogenetic tree and increase the variation in statistical annotations, thereby improving VQSR for XBS. XBS VQSR was run in SNP mode with a truth-set containing lineage and DR variants [17–19]. These variants reflect the diversity of the bacterium (Mtb lineages) and the entirety of the genome, enabling VQSR to build a model to identify variants in exactly such regions and as such avoid bias. The variants from the Goig et al. and reference dataset were converted to FASTA format, where positions represented by fewer than 95% of the samples were excluded. MTBseq and UVP were run in default mode.

IQ-TREE v2.1.2 was used to construct the Maximum Likelihood trees [20] which was plotted with Figtree v1.4.3 [21] and the resulting branch lengths were used to evaluate the potential presence of false positive variants.

## RESULTS

### Pipeline performance for analysis of WGS data from simulated culture isolates

At the highest coverage (100×) and with the low levels of contamination (≤5%), all pipeline approaches detected very few false positives and missed few variants, resulting in a 100% precision for SNPs and INDELs across the genome, except for UVP which obtained a slightly lower precision of 98% for SNP calling (Tables 1 and 2). Recall scores were highest for XBS and MTBseq-basic (97-99% for the SNPs and INDELs) compared to MTBseq-exrep and UVP which missed some variants (92% for SNPs and 91% for INDELs by MTBseq-exrep; 90% for SNPs and 89% for INDELs by UVP). The overall variant calling accuracy was highest for XBS and MTBseq-basic (F_1_ score 0.99 for SNPs, ≥0.98 for INDELs), somewhat lower for MTBseq-exrep (F_1_ score 0.96 for SNPs, 0.95 for INDELs), and lowest for UVP (F_1_ score 0.94 for SNPs and INDELs) (Figure 3). At 100× coverage MTBseq-basic identified an average 9.2% more true positive SNPs and 9.8% more INDELS per genome when compared to UVP (Table 1 and 2). XBS identified an average 9.0% more true positive SNPs and 8.1% more INDELs per genome when compared to UVP at 100× coverage.

**Figure 3:**
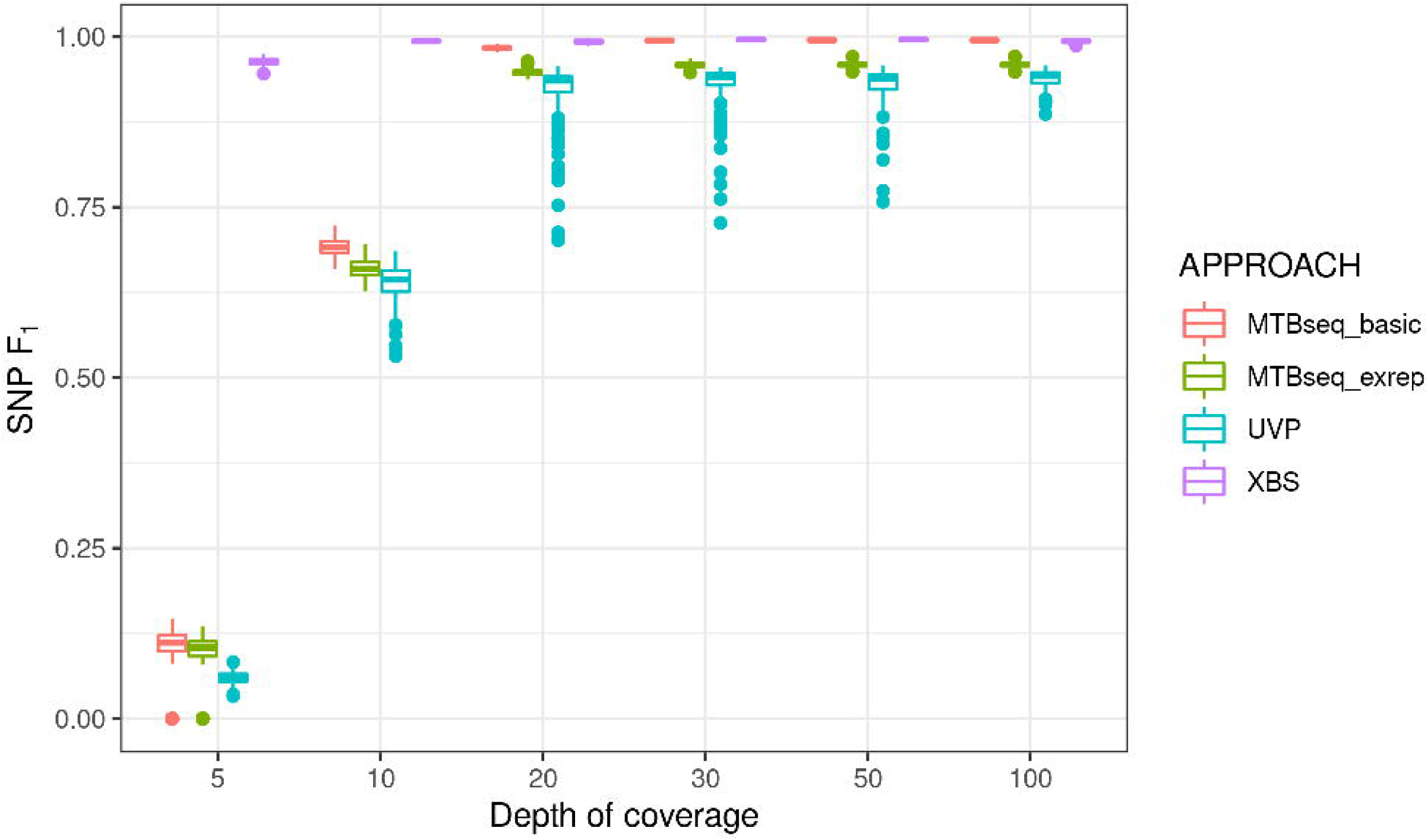
Performance (F_1_ scores) of four bioinformatic pipelines for SNP calling in simulated *Mtb* culture isolates at six levels of depth of *Mtb* genomic coverage.

Lowering the depth of *Mtb* genome coverage from 100× to 20× had minimal effect on accuracy scores. XBS’s precision and recal did not change and the F_1_ score deviated by ≤ 0.01 for SNPs and INDELs. The performance of MTBseq-basic and MTBseq-exrep also did not differ for these lower coverages with similar precision for SNPs and INDELs, a drop in recall by 2% for SNPs and 3% for INDELs, and the F_1_ score lowered by 0.01 for both SNPs and INDELs. UVP’s accuracy statistics did not change for INDELs, but precision, recall and F_1_ score for SNPs dropped slightly by 4%, 1% and 0.01, respectively.

At depths ranging from 20× to 100×, the performance for SNP and INDEL calling in the non-complex regions of the Mtb genome was high for all four pipelines (Figure 4 and Supplementary Table 1). Performance for variant calling in the complex regions was similar for MTBseq-basic and XBS, with an average SNP and INDEL precision of 100%, recall around 91% and the F_1_ around 0.95. Accuracy statistics for variant calling in the complex regions could not be calculate for UVP as complex regions are excluded from from its standard output. MTBseq-exrep’s exclusion of repetitive regions was less strict than UVP’s excluded loci and hence the former was able to identify a small number of variants in the complex regions.

**Figure 4:**
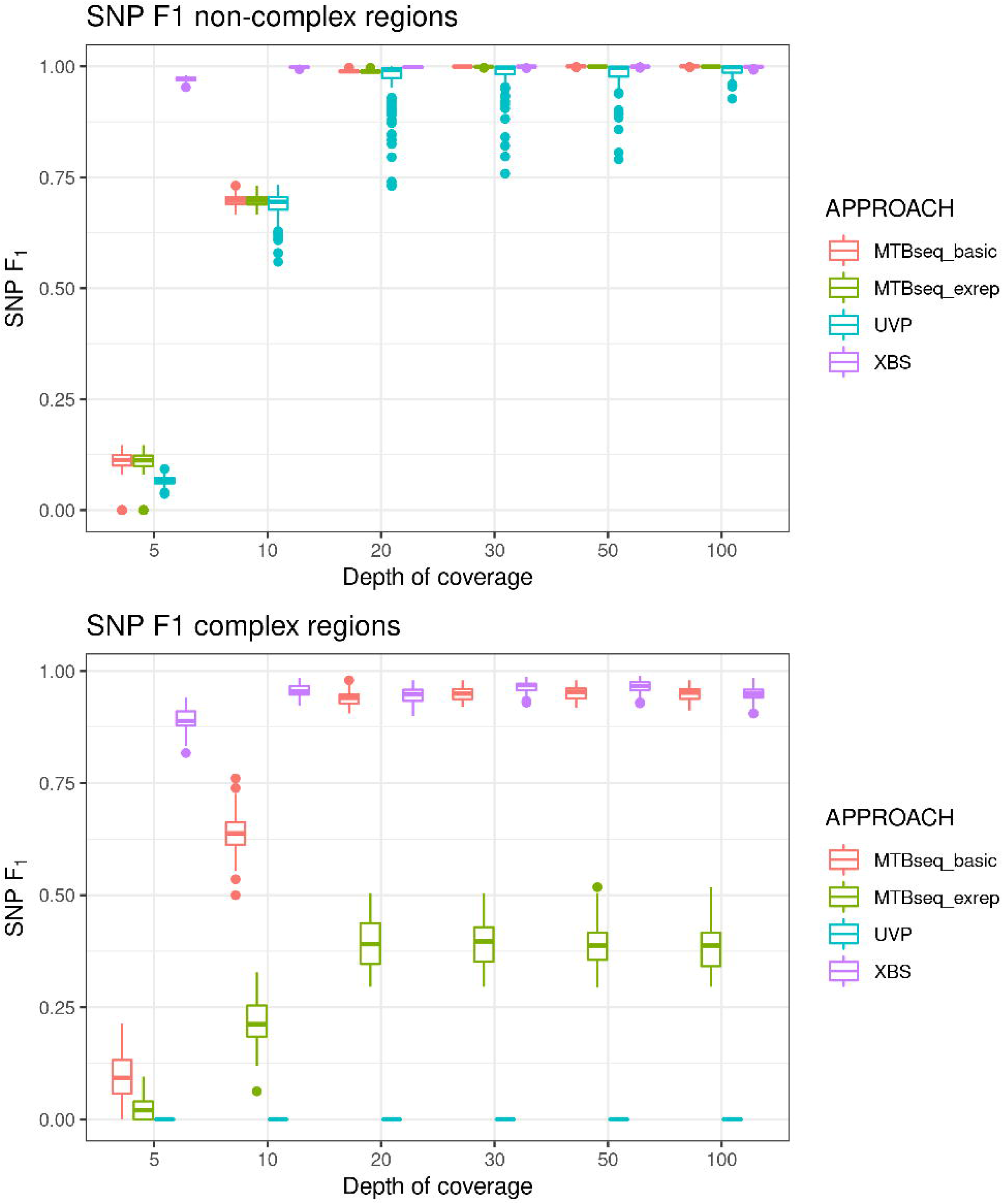
Performance (F_1_ scores) of four bioinformatic pipelines for SNP calling in simulated *Mtb* culture isolates at six levels of depth of *Mtb* genomic coverage, stratified by complex and non-complex regions of the genome.

At 10× depth of coverage, all four pipelines retained their precision but recall was affected. UVP and both MTBseq approaches missed many variants resulting in a recall of around 50% for SNPs and INDELs. Consequently, F_1_ scores dropped drastically for MTBseq-basic (0.69 for SNPs and 0.66 for INDELs), MTBseq-exrep (0.66 for SNPs and 0.63 for INDELs) and UVP (0.64 for both SNPs and INDELs). In contrast, XBS’s accuracy remained high, with recall at 99% for SNPs and 98% for INDELs and F_1_ scores of 0.99 for SNPs and INDELs (Figure 3). Only at an *Mtb* genome coverage of 5× was the performance of XBS noticeably affected, although performance remained largely accurate with F_1_ scores of 0.96 for SNPs and 0.95 for INDELs, a precision of 100% for both, and recall of 93% for SNPs and 91% for INDEL calling. XBS’s ability to call variants in both low coverage and complex regions was retained (Figure 4).

The type of low-level contaminant (*M. intracellulare, M. abscessus, M. avium, P. aeruginosa, S. epidermidis, H. sapiens*, NTM mixture or mixture of all 6 contaminants) only affected the F_1_ estimates of UVP due to false positive SNP calls when NTMs or *S. epidermidis* were present (Supplementary Figure 1 and Supplementary Table 1). The level of contamination (varying from 0-5%) also only affected UVP’s performance, with a decrease in precision and SNP F_1_ scores at higher levels of contamination (Supplementary Figure 2 and Supplementary Table 1).

### Pipeline performance for analysis of WGS data from simulated sputum samples

XBS outperformed both MTBseq approaches when analysing the data simulated to represent WGS directly from sputum. For SNPs, F_1_ scores ranged from 0.63-0.84 for XBS compared to 0.33-0.58 for MTBseq-basic and 0.31-0.59 for MTBseq-exrep. Using XBS, 49 to 77% of samples achieved F_1_ scores above 0.90, compared to 20 to 53% and 15 to 45% for MTBseq-basic and MTBseq-exrep, respectively. (Table 1 and 2). For INDELs F _1_ scores ranged from 0.61-0.81 for XBS, 0.32-0.63 for MTBseq-basic and 0.31-0.60 for MTBseq-exrep. Using XBS, 47 to 73 % of samples achieved F_1_ scores above 0.9, compared to 21 up to 58% and 16 up to 53% for MTBseq-basic and MTBseq-exrep, respectively. Plotting the theoretical depth of coverage against F_1_ score showed that XBS calls SNPs and INDELs with higher accuracy at low genomic depth of coverage compared to both MTBseq approaches (Figure 5).

**Figure 5:**
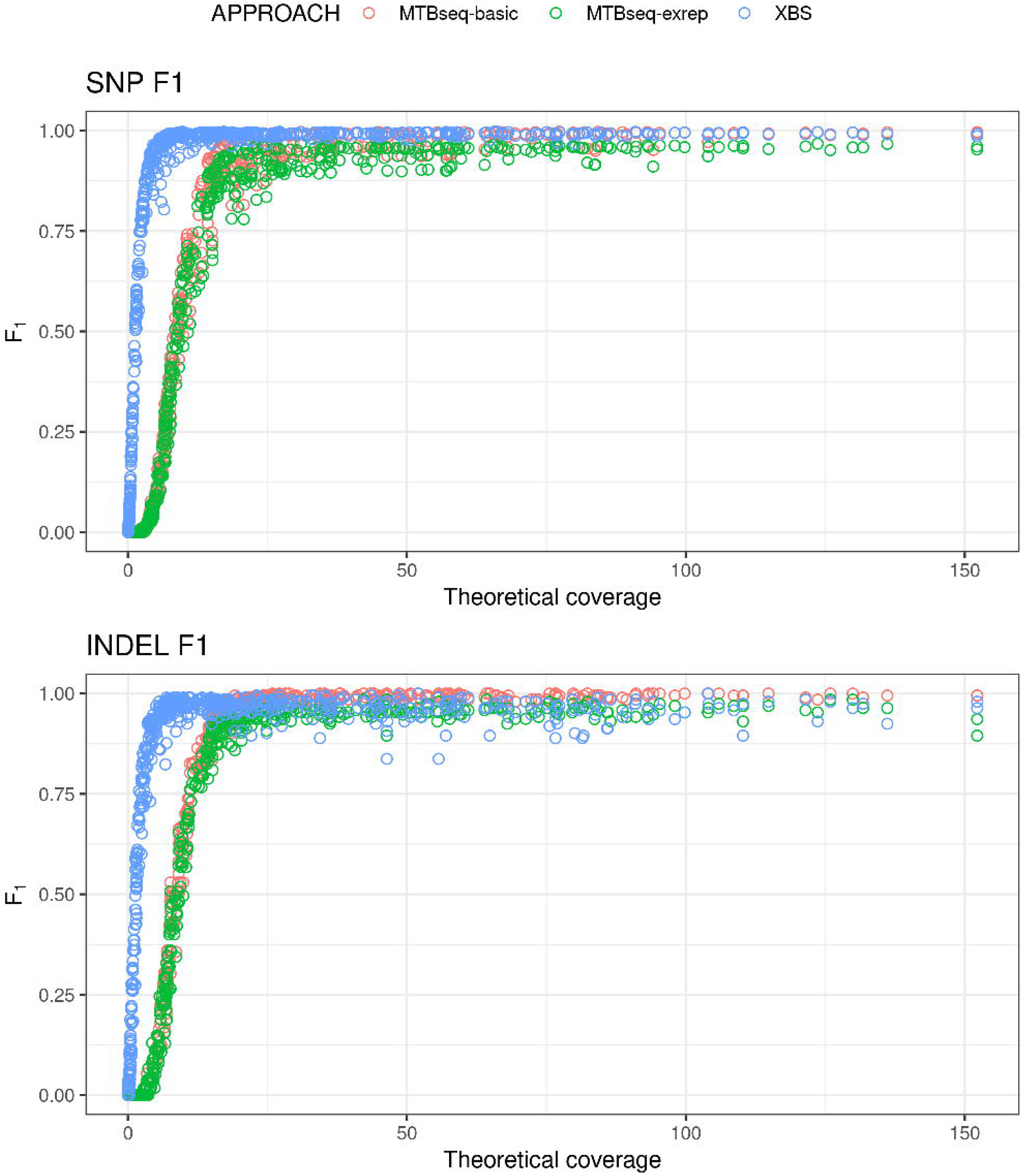
Performance (F_1_ scores) of different bioinformatic pipelines for SNP and INDEL calling from contaminated sputum samples at various levels of theoretical depth of *Mtb* genomic coverage.

SNP accuracy of XBS was unaffected by contamination level or type (Figure 6). In contrast, accuracy for MTBseq-basic and MTBseq-exrep depended on type and level of contamination. *H. sapiens* contamination did not affect the F_1_ score, *S. epidermidis* lowered the F_1_ score to 0.90 when the contamination level was ≥50%, and NTM and bacterial/human contamination mixtures reduced the F_1_ score when the contamination level was ≥75%. For INDELs, the MTBseq-basic pipeline performed slightly better than XBS when *Mtb* depth of coverage was ≥20×, with average F_1_ scores of 0.99 for MTBseq-basic, 0.95 for MTBseq-exrep and 0.96 for XBS respectively. (Supplemental Figure 3, Figure 5B).

**Figure 6:**
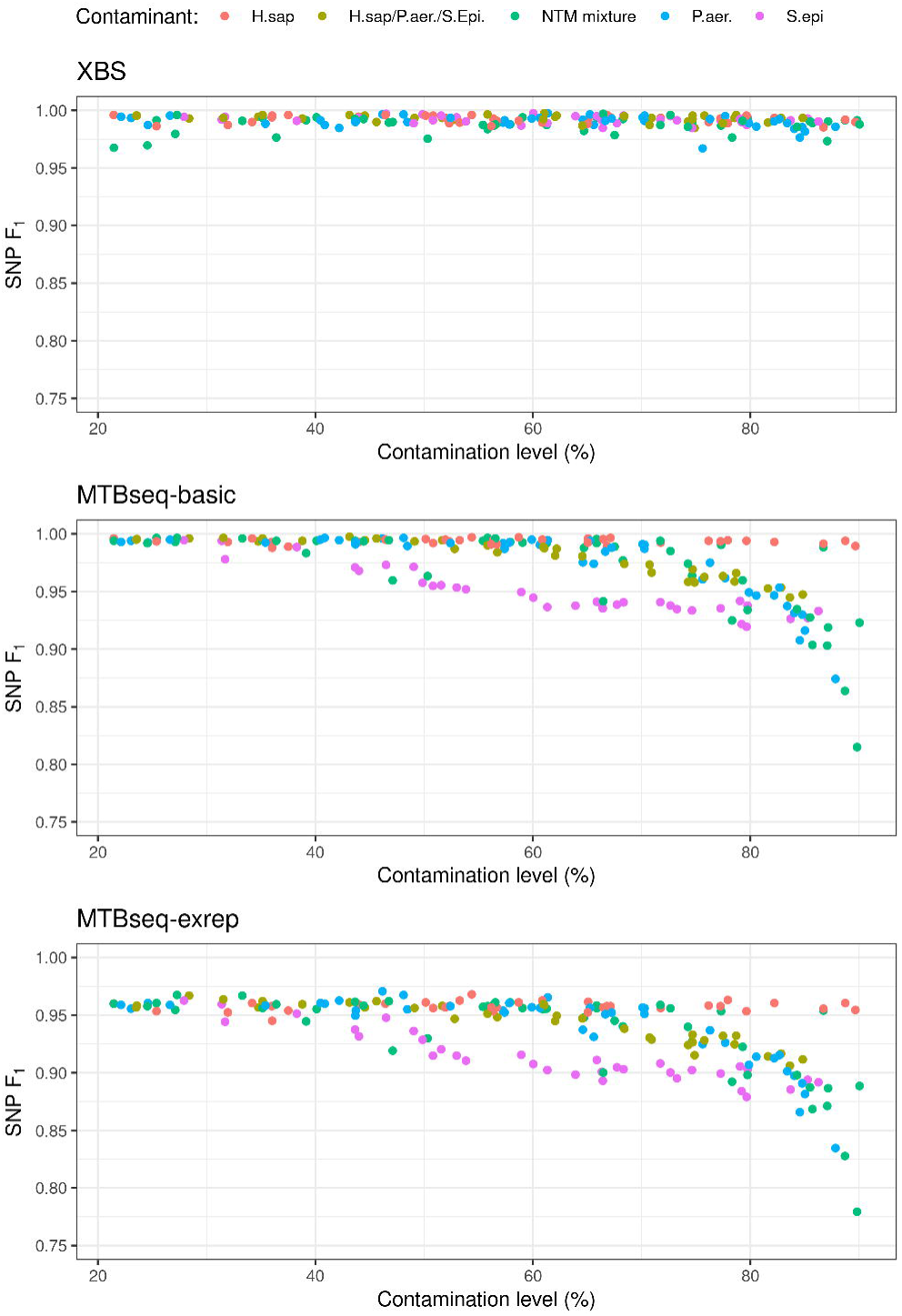
Performance (F_1_ scores) of three bioinformatic pipelines for SNP calling in sputum samples with minimum 20× Mtb coverage and various types and levels of contamination.

### Pipeline performance for analysis of WGS data from clinical culture isolates

MTBseq and XBS could analyse all samples from the Roetzer et al. dataset, whereas UVP excluded 33 of the 86 (38%) of samples. Of the 85 Sanger confirmed mutations, MTBseq-basic recovered 81, MTBseq-exrep 79, UVP 61 and XBS 84, corresponding to sensitivities of 95.3, 92.9, 71.8 and 98.8% respectively (Supplementary Table 3). The single variant missed by XBS was located right on the border of a repetitive region, resulting in reads with sub-optimal mapping qualities.

### Pipeline performance for analysis of WGS data from clinical sputum samples

UVP failed to analyse any sample included in the Goig et al. dataset as the contamination levels was above the 10% threshold for all samples. For the 10 Goig et al. samples and the 125 reference samples, XBS reported 11,977 variant positions (after exclusion of the ribosomal RNA regions), 13.9% more than the 10,514 variants reported by MTBseq. The number of variants called by MTBseq further reduced to 10,114 when variants within 12bp of each other were excluded.

There was no evidence of false positive variants when using XBS (no obvious branch extension for any sputum samples) with the highly conserved ribosomal RNA regions removed (Figure 7). When including the genes coding for ribosomal RNA obvious extended branch lengths were present for three samples (S02, S26 and S20, Supplemental Figure 4) due to VQSLOD scores for such variants that had fallen just within the within the positive VQSR model. When using MTBseq, there were also no obvious branch extensions but one sample (S26) showed a shorter branch length compared to its nearest-neighbours in the phylogenetic tree. This was the case for FASTA files in- and excluding SNPs within 12bp distance from each other (Supplementary Figures 5 and 6).

**Figure 7:**
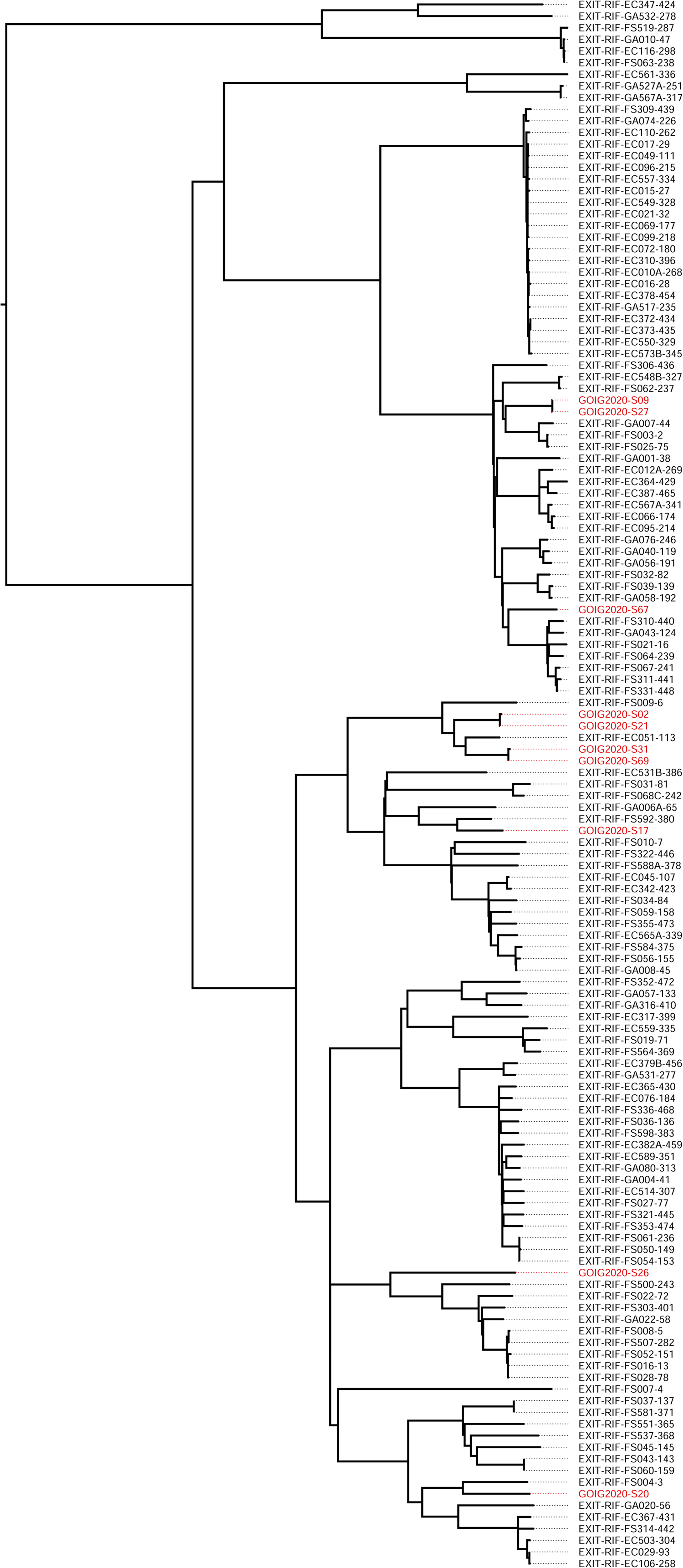
XBS Maximum Likelihood tree showing the location of Goig et al.’s sputum samples (marked in red) in relation to the reference dataset.

## DISCUSSION

We developed XBS and applied the joint variant calling and machine-learning-based variant filtering approaches, initially designed for human genome analyses, to a pipeline for Mtb WGS analyses. Using 1,200 simulated samples representing characteristics of WGS data from *Mtb* culture or directly from sputum samples, we demonstrated that XBS increases the performance in variant calling compared to existing pipelines (UVP and MTBseq), especially for WGS data from less-than-perfect, contaminated low *Mtb* burden samples. The strain simulation, variant calling and filtering approach presented here may also benefit the study of other bacteria where sequence coverage, complex genomes or contamination hinder accurate genetic variant identification.

We showed that current pipeline approaches perform well for SNP and INDEL calling when sequencing DNA extracts from decontaminated cultures with high (≥20×) depth, but accuracy decreases when depth of coverage is low (5-10×) or contamination levels are high (>50%). The novel XBS pipeline substantially outperformed other MTB pipelines for SNP and INDEL calling in *Mtb* at low coverage depth culture samples and highly contaminated, low coverage depth sputum samples.

When analysing WGS data from culture isolates at the current standard 30 to 100× depth coverage, all pipelines accurately called SNP and INDEL (F_1_ scores >0.90). Of the three pipelines assessed, UVP’s performance was slightly inferior given its lower precision (false positives variants) at higher (5%) contamination, particularly when the contaminant was an NTM. XBS and MTBseq-basic were not affected by low level (0-5%) contamination levels and identified on average 9% more SNPs and INDELs compared to UVP by investigating Mtb’s complex genomic regions. Identifying 9% more variants could greatly benefit transmission and genome-wide association studies.

At lower coverage (<20×), XBS was the only pipeline that could accurately call SNPs and INDELs, likely due to the joint calling and filtering processes that permit lower allele coverages. XBS’s accuracy remained high at 5× depth, where the modest drop in F_1_ score was due to a slightly lower recall rate and difficulty in accurately calling genuine INDELs due to coverage gaps. This is expected as, according to the Poisson distribution, only 99.3% of the genome is covered by at least one read at 5× coverage. The low-level contamination simulated for the culture samples did not affect the accuracy of the XBS or MTBseq pipelines. The UVP pipeline was however affected by both the level and the type of low-level contamination, such effects have been observed previously [22]. Considering these findings it is understandable that UVP uses a strict contamination cut-off, but the other pipelines show that variants can be identified more accurately despite the absence of such cut-offs. Sequencing at 5× depth using XBS resulted in average SNP F_1_ score of 0.96 (minimum 0.95) and average INDEL F_1_ scores of 0.95 (minimum 0.91), whereas the F_1_ scores of the other pipelines were ≤0.10 for both SNPs and INDELs. Such low-coverage sequencing could lower the costs by a factor of 10 compared to standard 50× coverage sequencing. In combination with low-cost library preparations, which is the main driver of sequencing cost, this could open the door to large-scale population sequencing projects in high TB-burden settings.

WGS data obtained directly from sputum is characterized by a low number of *Mtb* reads (theoretical coverage), a high the level of contamination, and presence of a mix of contaminants. The novel XBS pipeline showed superior performance for analysing such impure sequencing data. Due to the joint calling approach, XBS could analyse samples with much lower genomic coverage than the two MTBseq approaches. XBS successfully identified SNPs and INDELs in an average 73% of samples with 2,5 to 5 million paired-end reads, where MTBseq-basic only successfully analysed 50% and MTBseq-exrep 45% of such samples. By employing VQSR filtering, which identifies contaminant reads based on a multitude of statistical annotations, XBS’s performance was not affected by level or type of contaminants. Hard filtering, as is implemented in MTBseq-basic and -exrep, was not sufficient at high levels (>50%) of contamination because contaminant alleles may be interpreted as the most likely and therefore genuine allele, leading to false positives, once they reach coverage levels greater than the *Mtb* allele. For MTBseq, the type of contaminant affected the accuracy. High levels of human DNA did not affect accuracy as these are unlikely to map to the reference genome, but *S. epidermidis* contamination started to have an effect from 50% upwards as contaminant alleles then outnumber that of *Mtb*. The NTM-mix only affected accuracy at high contamination despite NTMs great genome similarity. This counterintuitive finding is likely because high levels of contamination are required before one of the NTM contaminants present in the mix approach 50% allele frequency.

Since it cannot be said exactly which samples underperform in terms of variant identification accuracy in a clinical dataset, it is best to ensure an acceptable minimum accuracy instead. When employing XBS on WGS data from real-life sputum, our data suggests that it may be prudent to restrict the analyses to those samples that present a coverage of ≥10×. A 10× cut-off would result in average SNP F_1_ scores of 0.99, minimum 0.98, and average INDEL F_1_ scores of 0.97, minimum 0.91, for 60% (60/100) of the samples in the 3,000,000 PE read dataset. MTBseq-basic would result in average SNP F_1_ scores of 0.91, minimum 0.53, and average INDEL F_1_ scores of 0.95, minimum 0.69, for the same samples.

Comparisons of the performance of Mtb pipelines is important but hampered by the absence of large datasets for which the true variants are known. To date, studies assessing the performance of *Mtb* pipelines have compared pipelines’ ability to identify transmission clusters as established through contact tracing or older molecular methods, or by comparing the detection of genomic drug resistance in relation to phenotypic tests or Sanger-confirmed variants [6,15,23,24]. These approaches suffer from important limitations. Contact tracing is complex and may not necessarily identify all clusters correctly [25]. Older molecular methods have significantly lower resolution than WGS so that all pipelines call clusters identified by these older methods with relative ease [26]. Using genomic drug resistance to compare pipelines is affected by the reference drug resistance mutation list used by each pipeline, whereas focussing on a limited number of Sanger-confirmed variants is not representative for the entire genome. The only other study that used simulated read datasets to compare combinations of mapper, caller and filtering methods found that the GATK variant caller in combination with VQSR consistently had the highest precision scores [27], supporting the findings of our study. This study however had multiple limitations. First, the GATK calling was performed for one sample at a time, which is not optimal for VQSR or low coverage samples. Second, for the VQSR truth sets, half of the samples’ variants with the best quality score were taken, an approach is problematic when high frequency contaminant alleles are present. Third, the use of the clinical CDC1551 strain prohibited accuracy assessment for the complex regions as the exact location of CDC1551 variants in relation to H37Rv cannot be established with certainty for complex regions.

We successfully overcame the limitations of prior *Mtb* pipeline accuracy studies by using an *in silico* approach to construct a fully representative variant truth set. The simulated dataset resembled clinical datasets by ensuring that some strains occurred once while others occurred several times. VQSR benefited from the presence of clonal strains as it improves the identification of variants in low coverage samples observed in other samples. Our datasets simulating culture isolates represented the range of depth (5 to 100×) and low levels of contamination (0-5%). Our simulated sputum datasets contained more than 21.4% contamination (mean 82.12%, maximum 99.99%) and thus very low levels of *Mtb* DNA, which correspond to the findings of a recent study of the clinical samples where most (51%) WGS data showed less than 5% *Mtb* DNA sequence reads [8]. The use of the simulated datasets allowed us not only to accurately quantify the performance of different pipelines for variant calling throughout the entire genome, including the complex regions, but also assess the effect of important characteristics that determine accuracy such as mycobacterial burden, level and type of contamination.

The excellent performance of XBS for the analysis of complex samples was confirmed when analysing WGS data obtained from clinical samples. The analyses of the WGS data from clinical culture isolates showed that XBS outperformed other pipelines (including pipelines not investigated in this study) in terms of sensitivity (Supplementary Table 3). The high specificity of XBS matches the findings of the culture simulations where the four pipeline approaches show similar recall scores. The analysis of WGS data obtained from DNA extracted directly from sputum samples confirmed that only XBS and MTBseq, but not UVP, could successfully analyse such data. While both pipeline showed high specificity (no evidence of false positive variants resulting in branch extension on phylogeny), the performance of XBS was superior to MTBseq as it allowed the identification of 13.9% more variants.

Several limitations remain to the novel XBS variant caller. First, XBS (and other pipelines) cannot analyse samples when NTM contaminant sequences exceed 20%. Such samples would require analyses by programs such as QuantTB that can potentially filter out NTM contaminants before Mtb variant identification as they resemble multiple infections [28]. Second, XBS requires multiple samples for each run. Previously inferred Genomic VCF’s can however be included from the Combine VCF step just before the VQSR, eliminating the need to batch new samples. Third, the highly conserved ribosomal RNA regions had to be excluded for optimal specificity as sequences in these regions from contaminating bacteria can map to the Mtb reference genome with very high confidence, making the variants in such regions indistinguishable in terms of the statistical annotations used by VQSR. Eliminating these regions may result in some loss of the genomic information, albeit small as this region represent only 0.1% of the genome Fourth, we used H37Rv as the reference genome for all analysis. If the Mtb ancestral genome would be used as the reference genome instead a VQSR truth-set could be constructed by aligning the ancestral genome to H37Rv and translating the latter its lineage and DR truth variants. When applying the XBS approach to other bacteria, one should employ well-established variants that occur throughout the entire genome for constructing the VQSR truth set. Finally, we were not able to compare the run time of XBS to the other pipelines as there were important differences in the analyses run other than the variant calling core and the number of samples analysed differed due to pipeline restrictions. However, elimination of the adapter removal and base recalibration steps reduces the overall processing time and exclusion of meta-genomic classification further reduces computing time. It also prevents the need for computing infrastructure with large memory requirements.

Direct-from-sputum WGS data contains a wealth of diverse bacterial contaminants besides that of human origin and all these contaminants occur at widely varying levels. It was not possible to represent this endless variety of bacterial contaminants in our simulation experiments hence the most commonly observed bacterial contaminants where used instead [2], NTMs were included because these pose a serious challenge for Mtb variant calling. Contaminant levels were simulated to match levels observed for genuine sputum samples [8]. As such it was possible to study the effect of the various contaminants and their levels on variant calling, this would not have been possible had all the endless contaminants observed for direct-from-sputum samples been used. To show that XBS was able to handle such genuinely diverse contamination two clinical WGS datasets were studied, one from cultured samples and one direct-from-sputum. Further studies are required to study the effect of the full diversity of contaminants that are observed for direct-from-sputum samples.

In conclusion, all pipelines studied (MTBseq, UVP, XBS) accurately analysed WGS data from *Mtb* culture isolates. Only XBS and MTBseq could accurately identify variants in low *Mtb* coverage and highly contaminated samples and XBS achieved higher performance parameters of all pipelines studies. High performance at low depth could decrease sequencing cost and improve WGS analysis directly from sputum samples. The accurate identification of variants in the complex genomic *Mtb* regions allow for improved resolution in transmission studies through increased genetic resolution and creates the ability to explore the functional role of variants in these complex regions. Taken together, the novel XBS pipeline sets the stage for the next generation of *Mtb* WGS studies.

## Authors and contributors

THH, RMW and AVR conceptualised the project and methodology. THH and LV curated the data and designed and scripted the software. RMW and AVR acquired the funding. THH, LV, RMW and AVR wrote, reviewed and edited the manuscript.

## Acknowledgements

We would like to thank the members of the Tuberculosis Omics ResearCH (TORCH) consortium for helpful discussions and particularly Elise De Vos for discussions surrounding sputum simulation. We thank the reviewers for helpful comments and Galo A. Goig et al. for providing additional direct-from-sputum sequencing data.

## Figures and tables

**Table 1: SNP calling accuracies across the entire genome for four Mtb pipelines**.

**Table 2: INDEL calling accuracies across the entire genome for four Mtb pipelines**.

