## Supplementary material for "Comprehensive and accurate genetic variant identification from contaminated and low coverage *Mycobacterium tuberculosis* whole genome sequencing data"

**Supplementary Table 1**

Can be found as part of this paper here: https://microbiology.figshare.com/

**Supplementary Table 2**

Can be found as part of this paper here: https://microbiology.figshare.com/

**Supplementary Table 3**

| Pipeline | Samples excluded (n=86) | Number correctly identified SNPs (n=85) | Sensitivity |
|---|---|---|---|
| MTBSEQ-basic | 0 | 81 | 95.3% |
| MTBSEQ-exrep | 0 | 79 | 92.9% |
| UVP | 33 | 61 | 71.8% |
| XBS | 0 | 84 | 98.8% |
| Anonymous A* | 0 | 79 | 92.9% |
| Anonymous B* | 0 | 62 | 72.9% |
| Anonymous C* | 17 | 75 | 88.2% |
| Anonymous D* | 0 | 77 | 90.6% |

* These numbers were reported by Walter et al. 2020

### Supplementary Figure 1

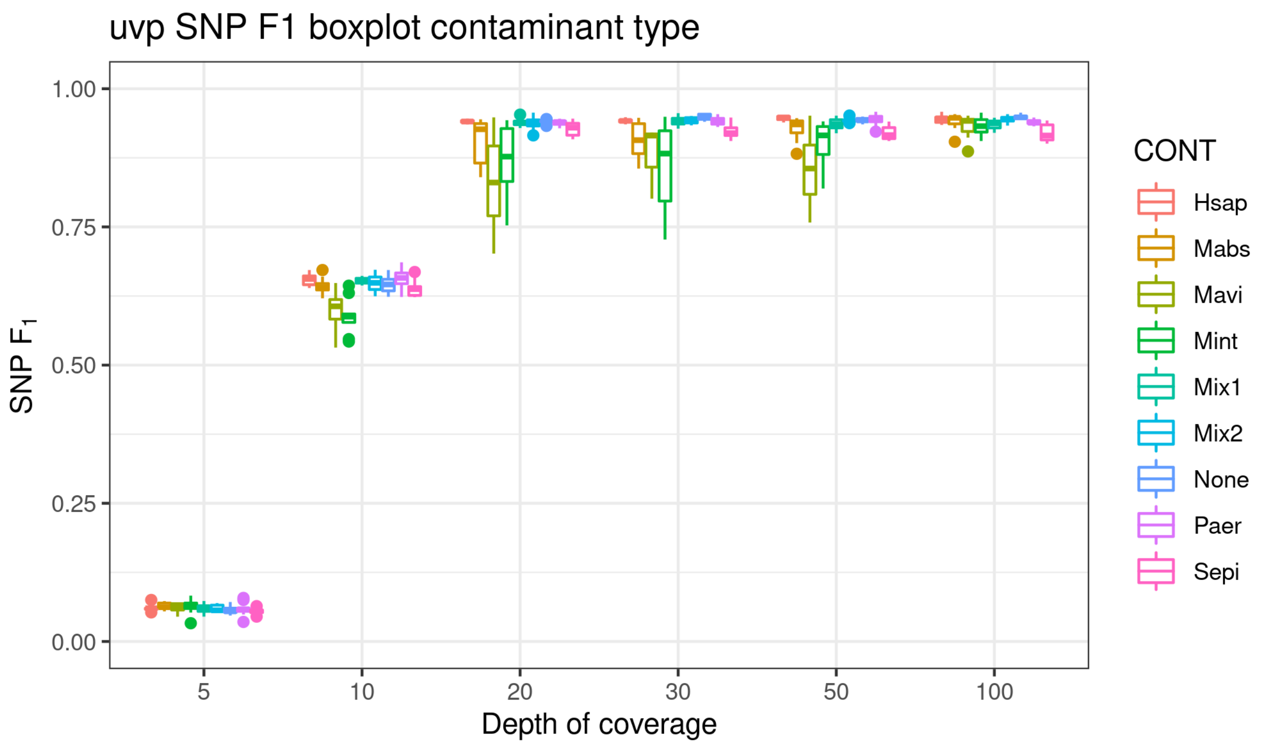


### Supplementary Figure 2

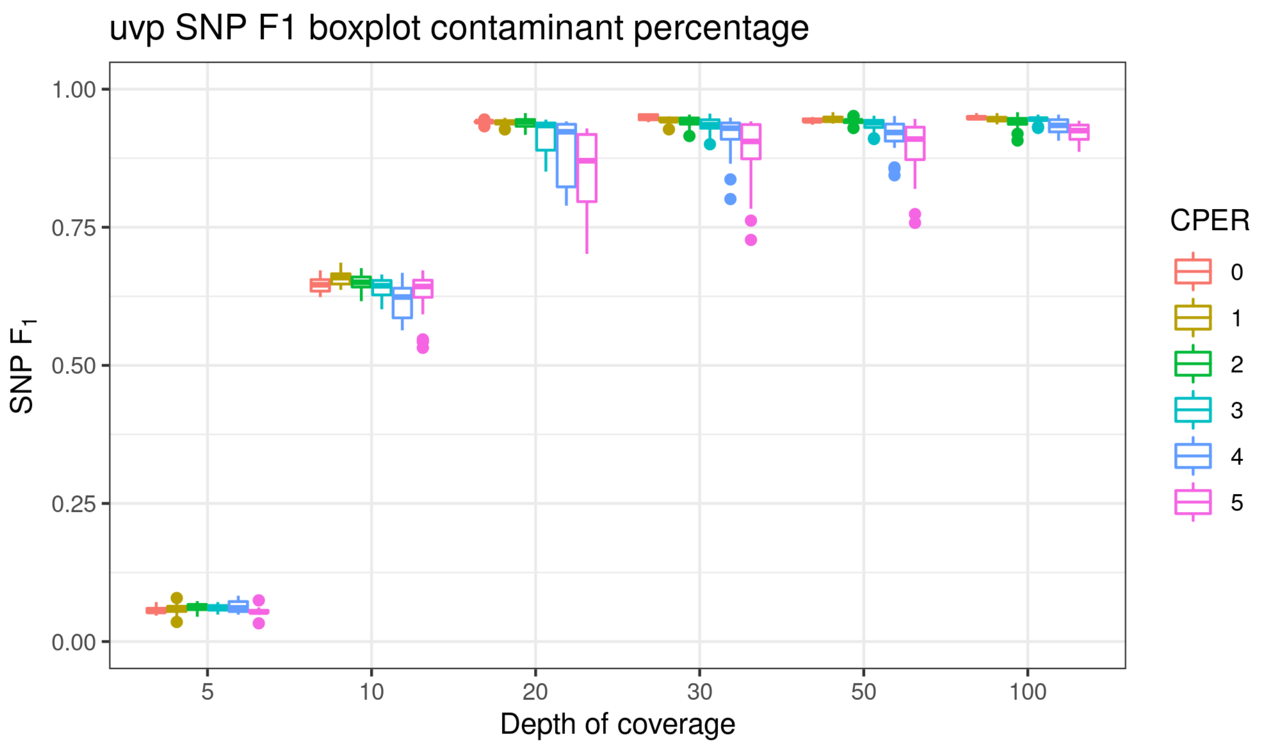

### Supplementary Figure 3

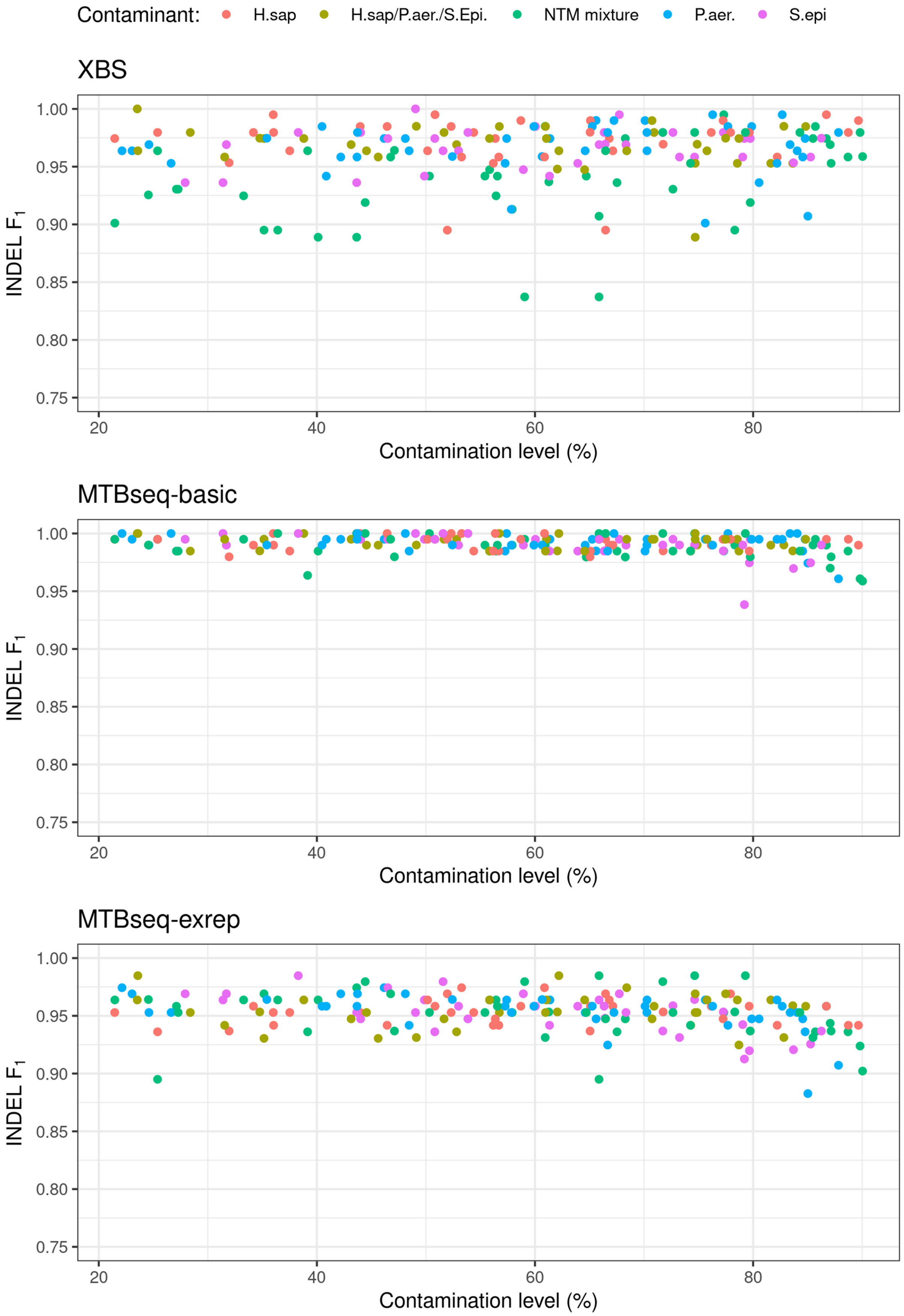

### Supplementary Figure 4: XBS including rRNA regions

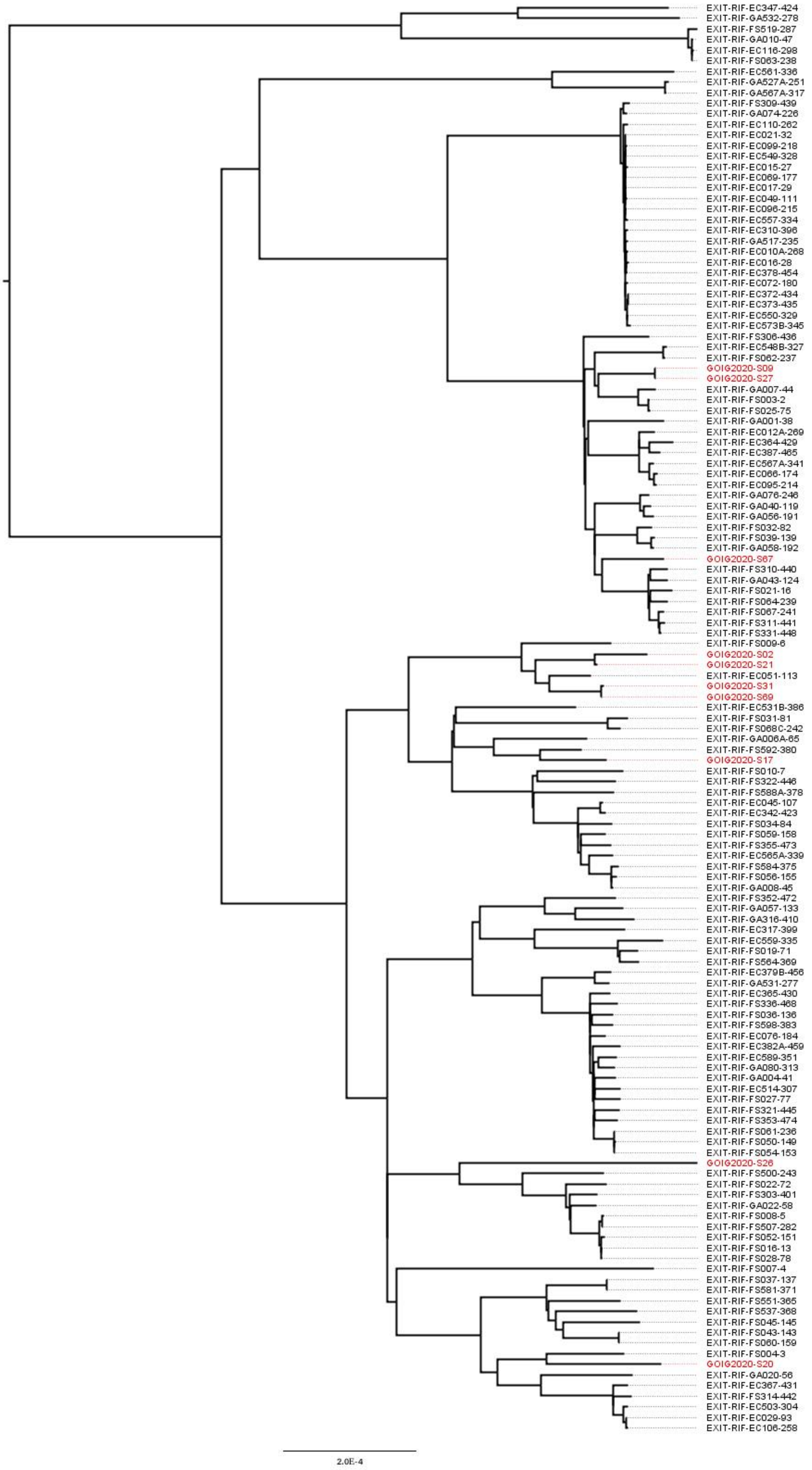

### Supplementary Figure 5: MTBseq including 12bp distance SNPs

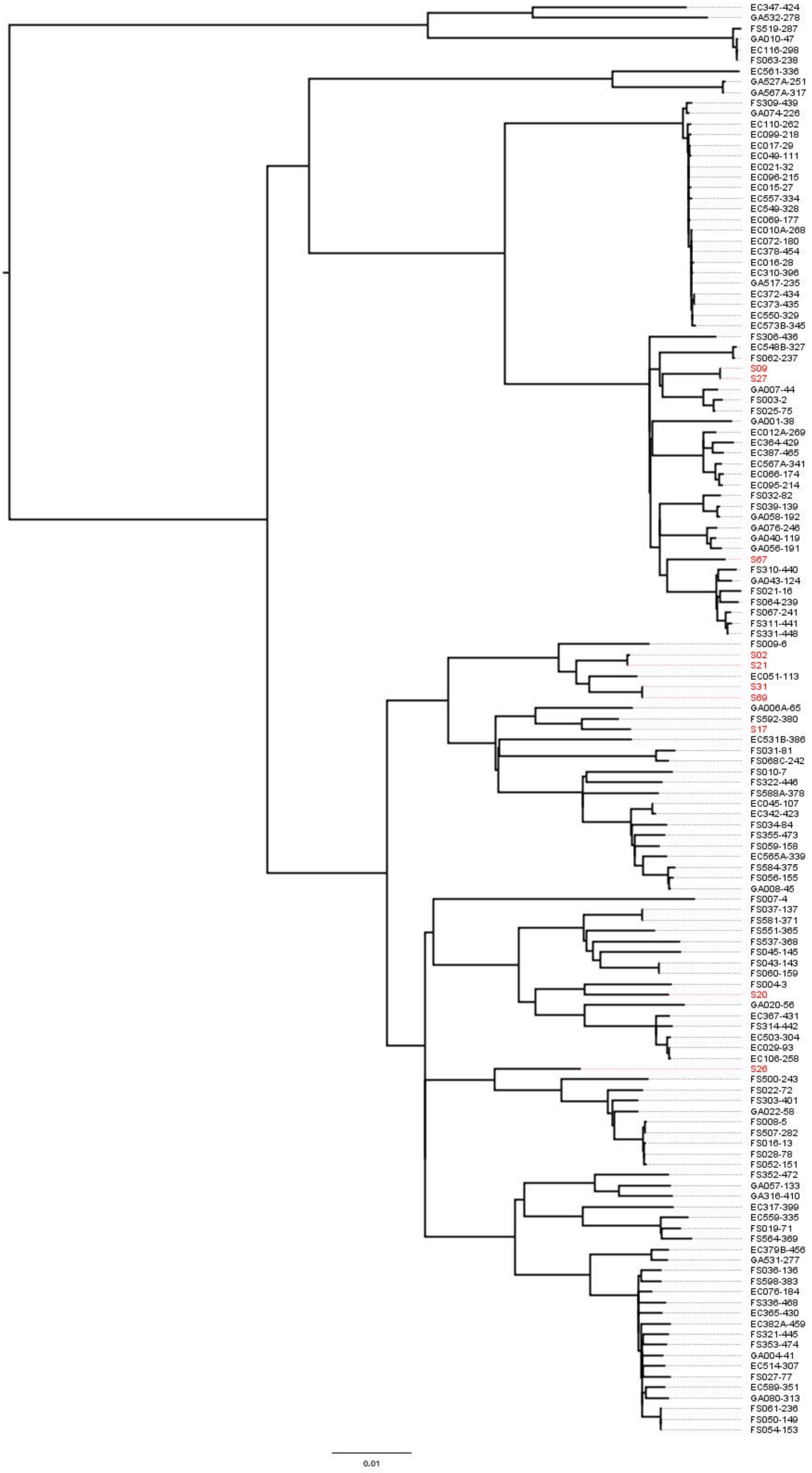

#### Supplementary Figure 6: MTBseq excluding 12bp distance SNPs

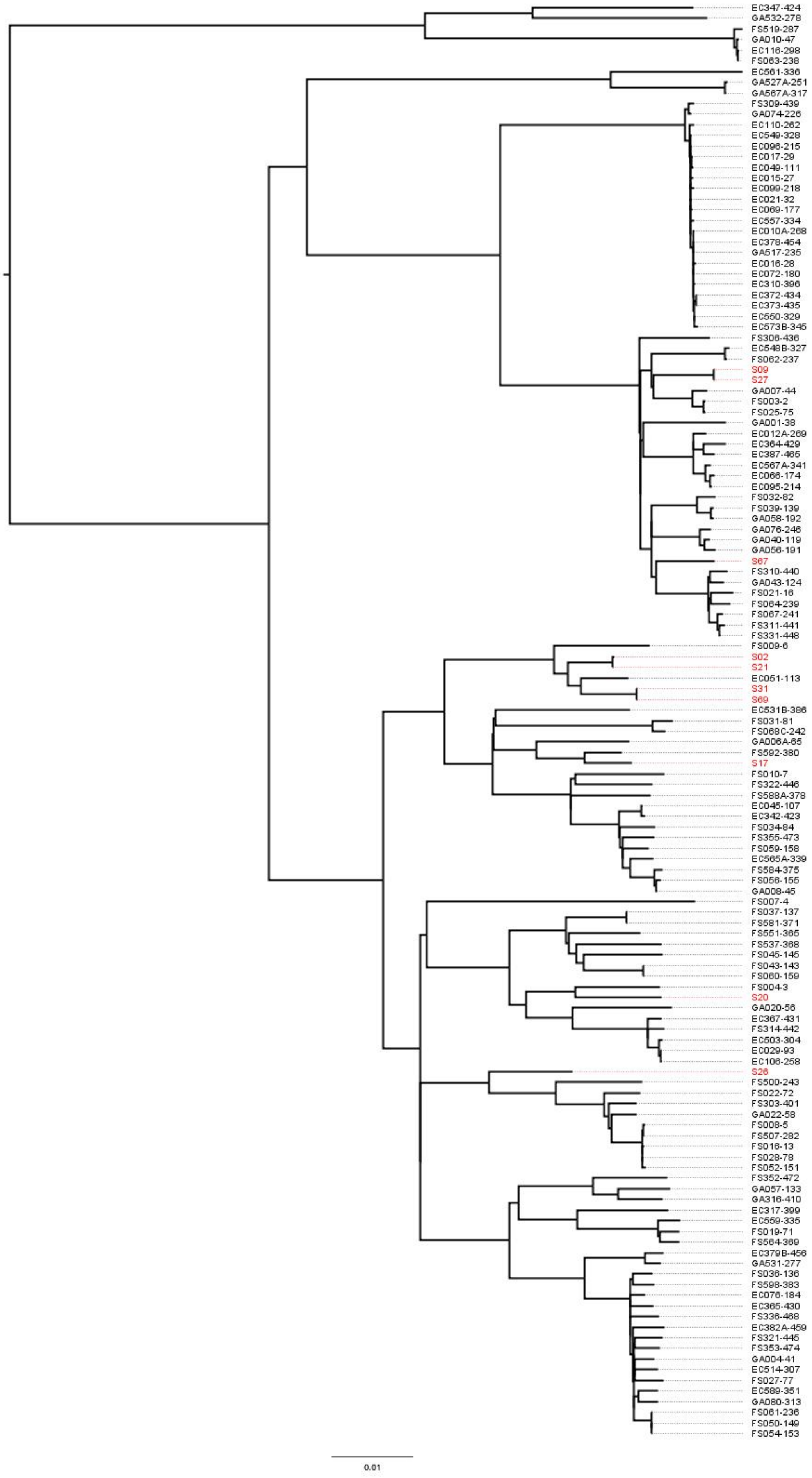
