## Supplementary material for "Comprehensive and accurate genetic variant identification from contaminated and low coverage *Mycobacterium tuberculosis* whole genome sequencing data": Table 2

|  | **XBS** | | | | **MTBseq-basic** | | | | **MTBseq-exrep** | | | | **UVP** | | | |
| --- | --- | --- | --- | --- | --- | --- | --- | --- | --- | --- | --- | --- | --- | --- | --- | --- |
| **Simulated dataset** | **PRECI-SION** | **RECALL** | **F_1_** | **F_1_ >0.90** | **PRECI-SION** | **RECALL** | **F_1_** | **F_1_ >0.90** | **PRECI-SION** | **RECALL** | **F_1_** | **F_1_ >0.90** | **PRECI-SION** | **RECALL** | **F_1_** | **F_1_ >0.90** |
| COV5 | 1.00 | 0.91 | 0.95 | 100 | 0.88 | 0.05 | 0.10 | 0 | 0.86 | 0.05 | 0.09 | 0 | 0.94 | 0.03 | 0.06 | 0 |
| COV10 | 1.00 | 0.98 | 0.99 | 100 | 1.00 | 0.50 | 0.66 | 0 | 1.00 | 0.46 | 0.63 | 0 | 1.00 | 0.48 | 0.64 | 0 |
| COV20 | 1.00 | 0.97 | 0.99 | 100 | 1.00 | 0.96 | 0.98 | 100 | 1.00 | 0.89 | 0.94 | 99 | 1.00 | 0.89 | 0.94 | 99 |
| COV30 | 1.00 | 0.97 | 0.98 | 100 | 1.00 | 0.99 | 0.99 | 100 | 1.00 | 0.91 | 0.95 | 99 | 1.00 | 0.89 | 0.94 | 99 |
| COV50 | 1.00 | 0.97 | 0.99 | 100 | 1.00 | 0.99 | 0.99 | 100 | 1.00 | 0.91 | 0.95 | 95 | 1.00 | 0.89 | 0.94 | 95 |
| COV100 | 1.00 | 0.97 | 0.98 | 100 | 1.00 | 0.99 | 0.99 | 100 | 1.00 | 0.91 | 0.95 | 98 | 1.00 | 0.89 | 0.94 | 98 |
| SPUTUM0500K | 0.73 | 0.57 | 0.61 | 47 | 0.46 | 0.28 | 0.32 | 21 | 0.46 | 0.27 | 0.31 | 16 | NA | NA | NA | NA |
| SPUTUM1000K | 0.88 | 0.70 | 0.74 | 67 | 0.56 | 0.36 | 0.40 | 28 | 0.55 | 0.34 | 0.38 | 27 | NA | NA | NA | NA |
| SPUTUM1500K | 0.86 | 0.67 | 0.73 | 57 | 0.55 | 0.45 | 0.48 | 41 | 0.55 | 0.42 | 0.46 | 40 | NA | NA | NA | NA |
| SPUTUM2000K | 0.87 | 0.68 | 0.74 | 62 | 0.64 | 0.52 | 0.55 | 45 | 0.64 | 0.48 | 0.53 | 40 | NA | NA | NA | NA |
| SPUTUM2500K | 0.91 | 0.75 | 0.79 | 70 | 0.70 | 0.61 | 0.63 | 58 | 0.70 | 0.57 | 0.60 | 53 | NA | NA | NA | NA |
| SPUTUM3000K | 0.86 | 0.79 | 0.81 | 73 | 0.69 | 0.58 | 0.61 | 47 | 0.69 | 0.54 | 0.59 | 45 | NA | NA | NA | NA |
